# Effects of tDCS Dosage on Working Memory in Healthy Participants

**DOI:** 10.1101/192419

**Authors:** Stevan Nikolin, Donel Martin, Colleen K. Loo, Tjeerd W. Boonstra

**Affiliations:** School of Psychiatry, University of New South Wales, Sydney, Australia; Black Dog Institute, Sydney, Australia; St. George Hospital, Sydney, Australia; Systems Neuroscience Group, QIMR Berghofer Medical Research Institute, Brisbane, Australia

## Abstract

**Background:** Transcranial direct current stimulation (tDCS) has been found to improve working memory (WM) performance in healthy participants following a single session. However, results are mixed and the overall effect size is small. Interpretation of these results is confounded by heterogeneous study designs, including differences in tDCS dose (current intensity) and sham conditions used.

**Aims:** We systematically investigated the effect of tDCS dose on working memory using behavioural and neurophysiological outcomes.

**Methods:** In a single-blind parallel group design, 100 participants were randomised across five groups to receive 15 minutes of bifrontal tDCS at different current intensities (2mA, 1mA, and three sham tDCS conditions at 0.034mA, 0.016mA, or 0mA). EEG activity was acquired while participants performed a WM task prior to, during, and following tDCS. Response time, accuracy and an event-related EEG component (P3) were evaluated.

**Results:** We found no significant differences in response time or performance accuracy between current intensities. The P3 amplitude was significantly lower in the 0mA condition compared to the 0.034mA, 1mA and 2mA tDCS conditions. Changes in WM accuracy were moderately correlated with changes in the P3 amplitude following tDCS compared to baseline levels (r = 0.34).

**Conclusions:** Working memory was not significantly altered by tDCS, regardless of dose. The P3 amplitude showed that stimulation at 1mA, 2mA and a sham condition (0.034mA) had biological effects, with the largest effect size for 1mA stimulation. These findings indicate higher sensitivity of neurophysiological outcomes to tDCS and suggests that sham stimulation previously considered inactive may alter neuronal function.

## Introduction

Transcranial direct current stimulation (tDCS), a form of non-invasive brain stimulation, is emerging as a promising technique for cognitive enhancement of mental processes such as learning, memory and attention (1, 2). Research suggests that augmentation of working memory in particular produces further benefits in other cognitive domains, including fluid intelligence (3, 4). Indeed, anodal tDCS to the left dorsolateral prefrontal cortex (DLPFC), a region of the brain involved in working memory processes (5), has been demonstrated to significantly improve performance in multiple systematic reviews (6-8). However, these reviews combined results from studies with heterogeneous stimulation parameters and experimental designs, resulting in mixed outcomes and an overall small sized effect. This heterogeneity complicates the identification of optimum parameters for working memory enhancement, such as tDCS dosage. Additionally, recent research has raised concerns that sham tDCS protocols, against which the effects of tDCS are typically compared, may provide a sufficient dose of electrical current to alter brain activity and produce behavioural effects (9, 10). Therefore, there is a need to systematically investigate tDCS dosage to understand the relative contribution of current intensity to working memory enhancement in healthy participants.

Although existing evidence suggests that tDCS to the DLPFC improves cognition (11), efforts to identify optimum tDCS dosage have produced contradictory findings. To date, only two studies have empirically examined the effect of varying tDCS current intensities on working memory outcomes in healthy participants (12, 13). Hoy et al. (13) investigated the impact of tDCS dosage (anodal 1mA and 2mA delivered to the left DLPFC) on working memory and found no significant differences in either response time or accuracy between conditions using the 3-back task. In contrast, Teo et al. (12) showed faster response times on the 3-back task with 2mA tDCS, but not 1mA. Dedoncker et al. (8) conducted a meta-analysis of the cognitive effects of tDCS and found that current intensity did not modulate accuracy or response times. The authors concluded that significant heterogeneity between trials may have reduced the ability to assess predictors of response, including the effect of tDCS dose. Therefore, there remains a need to test the effect of tDCS dose on working memory outcomes within a single trial.

The effects of tDCS dose are also relevant for sham protocols used as control conditions. Sham tDCS involves a short (10-60s) ramping up of current to a level similar to active tDCS conditions, followed by a ramp down phase of similar duration (14). This variation in current intensity replicates the sensation of active tDCS by generating paraesthetic effects, thus preserving blinding and allowing for a comparison of tDCS outcomes against placebo effects (15). Differences exist between studies regarding the precise duration of current ramping (15-18), and the background level of current output following the ramp down phase (19, 20). Sham protocols are commonly thought to be neurologically inert due to the low levels of current used (21), however, recent findings tentatively suggest that sham tDCS may alter neural activity. Boonstra et al. (9) collected resting-state electroencephalography (EEG) data before and after tDCS and found a similar generalised reduction in mean brain frequency in both active and sham conditions, differing only in effect size. Likewise, a multicentre trial of tDCS for the treatment of depression found that participants receiving sham stimulation had greater improvement in mood during the sham-controlled phase of the experiment compared to 2.5mA of tDCS (10). Similar responses to sham stimulation have been observed across various treatments and patient populations (22, 23), including major depressive disorder (24-26), but to date have been attributed to the placebo effect. If the low dosage of tDCS used in sham protocols is biologically active, the implications could be far reaching. This is particularly relevant in the context of therapeutic tDCS interventions, which estimate tDCS efficacy against these sham protocols.

Physiological measures can be used to objectively quantify the modulatory effects of tDCS and assess the effects of differing stimulation intensities. Changes in motor cortex excitability following tDCS, assessed using transcranial magnetic stimulation (TMS) induced evoked potentials, suggest that lower intensities (0.5 – 1mA) produce greater effects in healthy participants (27). However, these findings have not been consistently replicated (28-30), and it is unclear whether these results can be generalised to the DLPFC. Neurophysiological measures derived from electroencephalography (EEG) can supplement behavioural measures (31), and may be more sensitive to tDCS-induced changes in working memory than behavioural outcomes alone (32). Specifically, analysis of frontal activity during a working memory task using event related potentials (ERPs) has shown increases in a component (P3) thought to be generated by the DLPFC (33). These measures can hence be used to test whether different stimulation intensities are biologically active.

### Aims

The current study empirically tested the dose-response relationship of current intensity and working memory performance in healthy participants using behavioural and neurophysiological measures. Higher current intensities were hypothesised to result in greater cognitive enhancement based on prior studies (12). Two typical sham stimulation protocols were also assessed, in addition to a control condition which involved no stimulation, to test whether sham stimulation is biologically active.

## Methods

### Participants

All participants gave written informed consent prior to participation and the experimental protocol was overseen by the University of New South Wales (UNSW) Human Research Ethics Committee (HC13278). Participants were recruited through advertising on a UNSW website and were thus predominantly university students (age: 22.9 ± 4.3 years). Exclusion criteria included significant psychological or neurological illness, excessive alcohol or illicit substance abuse, smoking, and ambidextrous or left-handed applicants assessed using the Edinburgh handedness test (34). One hundred participants in total were allocated equally to five conditions using a parallel group, single blind, study design such that there were twenty participants per condition. Stratified randomisation was used to allocate participants to each condition based on baseline working memory performance.

### Procedure

The study consisted of two experiments:

#### Experiment 1

Initially, 40 participants were randomly allocated to receive either 2mA tDCS or sham tDCS (‘Sham1’). Participants completed a working memory task, followed by EEG setup lasting approximately 15-20 minutes and then resting state EEG (Figure 1). Preliminary analysis of this data prompted a second experiment using additional current intensities to better assess the role of tDCS dosage and examine the potential effect of sham tDCS.

**Figure 1.**
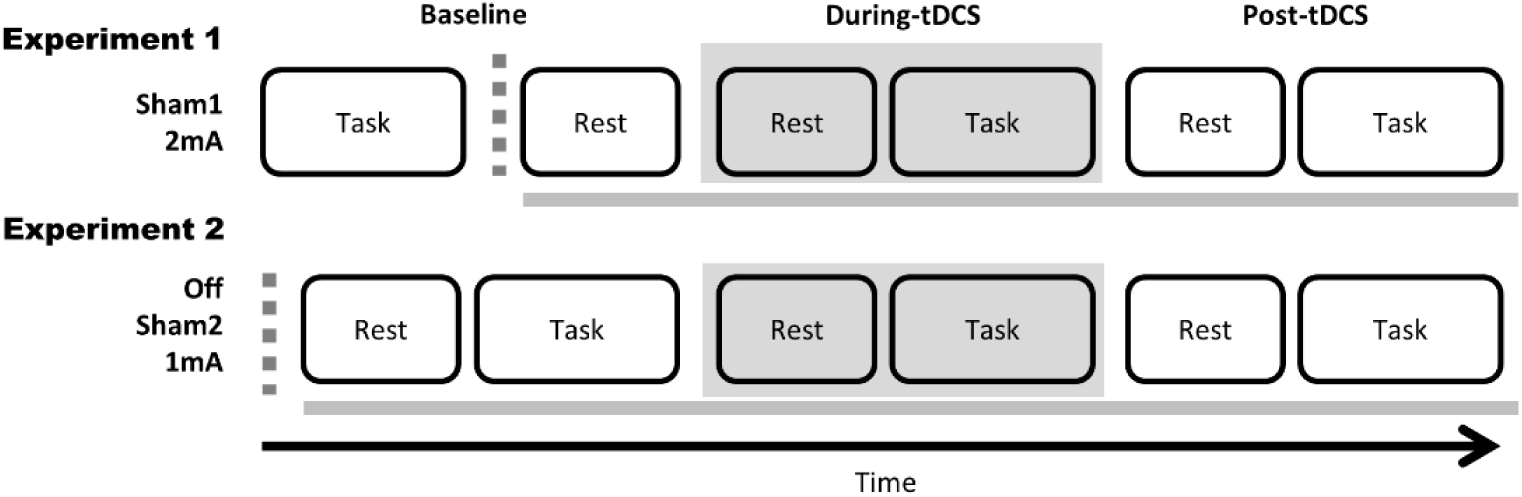
Study protocol. Recruitment occurred in two experiments; in Experiment 1 participants were allocated to receive either Sham1 or 2mA tDCS; in Experiment 2 participants were allocated to receive either Sham2, 1mA tDCS, or no stimulation at all in an ‘Off’ condition in which the electrode leads were left unplugged for the entirety of the experiment. The dotted grey line represents when EEG setup occurred and took approximately 15-20 minutes. The solid grey line shows when EEG was collected.

#### Experiment 2

60 participants were similarly randomised to one of three conditions; an ‘Off’ condition with the electrode leads left unplugged, a second sham condition using a different device (‘Sham2’), and 1mA tDCS. A modification was made to the order of events in the study design to capture baseline task-related EEG activity. Therefore, EEG setup was done first, followed by resting state EEG and then the working memory task (Figure 1).

### Transcranial Direct Current Stimulation

Stimulation was given for 15 minutes using 4cm × 4cm electrodes (16cm^2^) with the anode placed on the left DLPFC (F3 according to the International 10-20 EEG system) and the cathode on the right DLPFC (F4). This resulted in a current density over the electrode area of approximately twice that produced using standard 5cm × 7cm electrodes (35cm^2^), which has been associated with greater improvements to cognitive performance following a single session of tDCS (8). For both 1mA and 2mA tDCS conditions, stimulation was delivered using an Eldith DC-stimulator (NeuroConn GmbH, Germany). Sham tDCS was delivered using either an Eldith DC-stimulator (Sham1), or a tDCS-CT stimulator (Sham2; Soterix Medical Inc., New York). Sham1 stimulation ramped up to 2mA over 30 seconds, remained at that level for 30 seconds, ramped back down over 30 seconds, and then the machine default operation during the off-stimulation mode produced a constant background current of 0.016mA (verified using an independent ammeter). The parameters for Sham2 were pre-programmed for a clinical trial (35), and consisted of a 10 second ramp up to 1mA followed immediately by a ramp down over 60 seconds, and then the machine default operation during the off-stimulation mode produced a constant background current of 0.034mA. The total amount of charge (millicoulombs) delivered during the experiment session for each stimulation condition was: Off-0mC; Sham1-134mC; Sham2-66mC; 1mA-930mC; 2mA-1860mC.

### Tasks

The visual 3-back working memory task, adapted from Mull et al. (36), was administered using Inquisit 4 software (Version 4, Millisecond Software). This task required participants to press a key (spacebar) when a displayed letter matched a letter shown three trials previously. Working memory performance was assessed using d-prime, a measure of discriminate sensitivity (37), and response times for correct responses only. Participants were given the opportunity to practice the 3-back task for 5 minutes under the observation of the experimenter to ensure that they understood task instructions.

Previous tDCS studies examining cognition used crossover designs to minimise the impact of inter-individual variability. However, given the number of conditions and task presentations investigated in the present study, a parallel design was selected instead to minimise the risk of carryover effects (i.e. practice effects on the task). To account for individual differences in baseline performance levels, participants completed the 3-back task and were then stratified according to d-prime performance scores into the following tertiles: Low = 1.5-2.5, Mid: 2.5-3.5, High: >3.5. Cut-offs for stratification were based on data obtained from a previous study of tDCS and working memory (38).

### EEG Data Acquisition

EEG data was acquired using a TMSi Refa amplifier (TMS International, Oldenzaal, Netherlands). A 33-channel head cap with water-based electrodes was used for 31 EEG recording channels (Figure 2). Sites F3 and F4 were reserved for tDCS-electrode channels.

**Figure 2.**
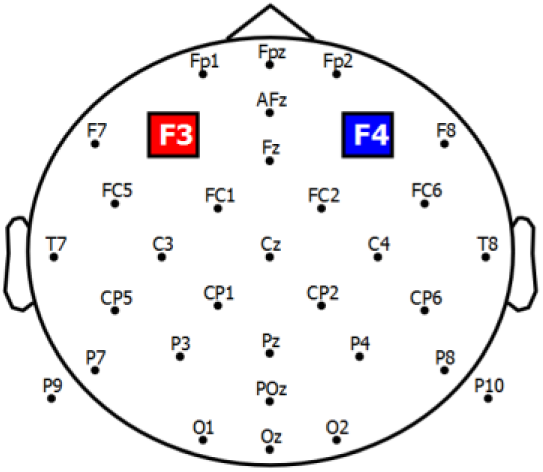
EEG and tDCS electrodes. A total of 31 EEG electrode channels were used from the following sites: O1, Oz, O2, P4, P3, Cz, Poz, Pz, P7, P9, Fc5, Fc1, T7, C3, Cp5, Cp1, P8, P10, Fc6, Fc2, T8, C4, Cp6, Cp2, Fp1, Fpz, Fp2, F8, F7, Fz, and Afz. tDCS electrodes (4cm × 4cm) were placed with the anode on the left DLPFC (F3) and the cathode on the right DLPFC (F4).

EEG analysis was conducted using custom-developed Matlab scripts (v.R2016a; MathWorks) in addition to the Fieldtrip toolbox (39). EEG data was sampled at 1024Hz and filtered using both a bandpass filter (0.5-70 Hz) and a notch filter at 50Hz to remove line noise. Data were inspected using a semi-automated algorithm to remove epochs containing artefacts. Independent components analysis was then used to remove eye blink and muscle artefacts.

Event related potentials (ERPs) were calculated using all stimuli from the 3-back task (including both targets and non-targets) and baseline corrected using the mean amplitude from 200ms to 500ms prior to stimulus onset. Previous studies using similar working memory tasks have identified P3 as a component of interest, with results typically obtained from frontal midline channels (Fz) (13, 33). Therefore, the mean amplitude of the P3 component was extracted by averaging ERPs from the Fz electrode channel in the 220-420ms time window following stimulus onset. Topographies of the P3 component were created for each stimulation condition to observe differences in the spatial distribution and magnitude of the evoked brain response.

### Statistical Analysis

Analyses were conducted using SPSS-software (SPSS 22.0 for Windows; SPSS Inc.). Baseline data were assessed for differences between conditions using one-way ANOVAs and chi-squared tests for continuous and categorical variables, respectively. These included demographic characteristics (age, gender and education level), as well as baseline performance on the working memory task and P3 amplitude.

Mixed effects repeated measures model (MRMM) analyses were conducted for both working memory and neurophysiological outcomes. Scores greater than three standard deviations from the mean were considered outliers and excluded from analyses. Residuals were inspected following each analysis for skew and kurtosis to ensure adequate convergence of model parameters. Planned contrasts were conducted comparing the Off condition to Sham1 and Sham2 to determine whether sham stimulation was significantly different from no stimulation at all. Hedges’ *g* was used to calculate effect sizes using differences in means divided by the pooled standard deviation.

For examining working memory outcomes, factors included were Time (baseline, during-tDCS, and post-tDCS), Condition (Off, Sham1, Sham2, 1mA, and 2mA), Baseline Performance (Low, Mid, and High tertiles from d-prime stratification), and the Time × Condition interaction. Participants were included as a random factor.

For the ERP analyses, only baseline and post-tDCS data were used due to the presence of EEG artefacts during tDCS. Additionally, due to the difference in task presentation order between experiments 1 and 2, baseline EEG recordings during the 3-back task were only available for Off, Sham2, and 1mA tDCS conditions. Therefore, ERP analyses were done in two stages. In the first stage, factors for MRMM analysis consisted of Time (baseline and post-tDCS), Condition (Off, Sham2, and 1mA), and Time × Condition. Participants were included as a random factor. Data were available for all conditions at the post-tDCS timepoint only, therefore, in the second stage of analysis these were compared using a one-way ANOVA with just the factor of Condition (Off, Sham1, Sham2, 1mA, and 2mA).

To confirm that behavioural and neurophysiological measures examined similar facets of working memory functioning, Pearson correlations were used to test the relationship between changes in working memory performance (both response times and d-prime) and P3 amplitude. Change scores were calculated by subtracting Baseline scores from Post scores. Correlations were conducted using combined data from participants in Off, Sham2, and 1mA conditions (for whom there was baseline neurophysiological data).

Lastly, blinding to tDCS condition was tested using a Pearson chi-square test examining participant guesses. Post-hoc testing of adjusted standardised residuals was used to determine whether participant guesses were more accurate for certain conditions over others. Residuals greater than an absolute value of 1.96 were considered significant (this represents the boundary for a 95% confidence interval).

## Results

One participant allocated to the Off condition discontinued the experiment in the post-tDCS period due to a strong sensation of nausea. Thus, the sample size in the Off condition was reduced to 19. Adverse events for all conditions are presented in Supplementary Table 1. The experimental protocol was otherwise well tolerated, with no significant differences in adverse event proportions among the five conditions except for skin redness which only occurred in active conditions (i.e. following 1mA and 2mA stimulation).

**Table 1.** Baseline comparison of participant characteristics. Significance was calculated using Pearson’s chi-squared test for categorical variables, and one-way ANOVA for continuous variables. Values represent: mean (standard deviation). ERP: event related potential. \**p*<.05

|  | Off | Sham1 | Sham2 | 1mA | 2mA | *p*-value |
| --- | --- | --- | --- | --- | --- | --- |
| Sample (n) | 19 | 20 | 20 | 20 | 20 | - |
| **Demographic details** |  |  |  |  |  |  |
| Age | 21.9 (2.9) | 22.9 (3.1) | 21.9 (3.3) | 22.8 (3.1) | 25.1 (7.0) | 0.123 |
| Education (years) | 15.1 (2.0) | 15.8 (2.3) | 14.7 (2.1) | 15.6 (2.1) | 16.4 (2.6) | 0.161 |
| Gender (F/M) | 11/8 | 8/12 | 9/11 | 12/8 | 12/8 | 0.587 |
| **Working Memory** |  |  |  |  |  |  |
| D-prime | 2.47 (0.72) | 2.58 (0.69) | 2.58 (0.66) | 2.48 (0.50) | 2.63 (0.84) | 0.937 |
| Response time (ms) | 663 (163) | 725 (163) | 710 (164) | 728 (146) | 709 (193) | 0.751 |
| **ERP<sup>1</sup>** |  |  |  |  |  |  |
| P3 mean amplitude | -1.07 (1.13) | - | -0.50 (0.97) | -0.78 (1.53) | - | 0.369 |

**Table 2.** Participant blinding. Adequacy of blinding was assessed by asking participants to guess whether they were in a sham condition (i.e. Off, Sham1, or Sham2) or an active condition (i.e. 1mA, 2mA) at the end of the experiment. Significance was calculated using a Pearson chi-square test. Z-scores were obtained using adjusted standardised residual scores. \**p*<0.05

|  | Off | Sham1 | Sham2 | 1mA | 2mA | *p*-value |
| --- | --- | --- | --- | --- | --- | --- |
| Participant Guess |  |  |  |  |  |  |
| sham tDCS | 12 | 6 | 5 | 4 | 8 | 0.043* |
| active tDCS | 7 | 14 | 15 | 16 | 12 |  |
| Adjusted Residuals |  |  |  |  |  |  |
| z-score | 2.8* | -0.6 | -1.1 | -1.6 | 0.5 |  |

Demographic information such as age, gender and years of education were well balanced between groups (see Table 1). Similarly, there were no baseline differences for working memory performance and P3 amplitude.

### Working memory performance

For 3-back d-prime scores there were significant main effects of Time (*F*_(2,131.7)_ = 10.5, *p* < 0.001), and Baseline Performance (*F*_(2,111.9)_ = 113.9, *p* < 0.001), but no main effect of Condition (*F*_(4,92.2)_= 1.00, *p* = 0.414). The Time × Condition interaction was not significant *(F*_(8,131.7)_ = 1.34, *p* = 0.228); see Figure 3A and 3B for raw scores and plots of estimated marginal means, respectively. Planned simple contrasts for the post-tDCS time point revealed no significant effect of Off compared to Sham1 (*p* = 0.558, Hedges’ *g* = 0.12) or Sham2 (*p* = 0.426, Hedges’ *g* = −0.50).

**Figure 3.**
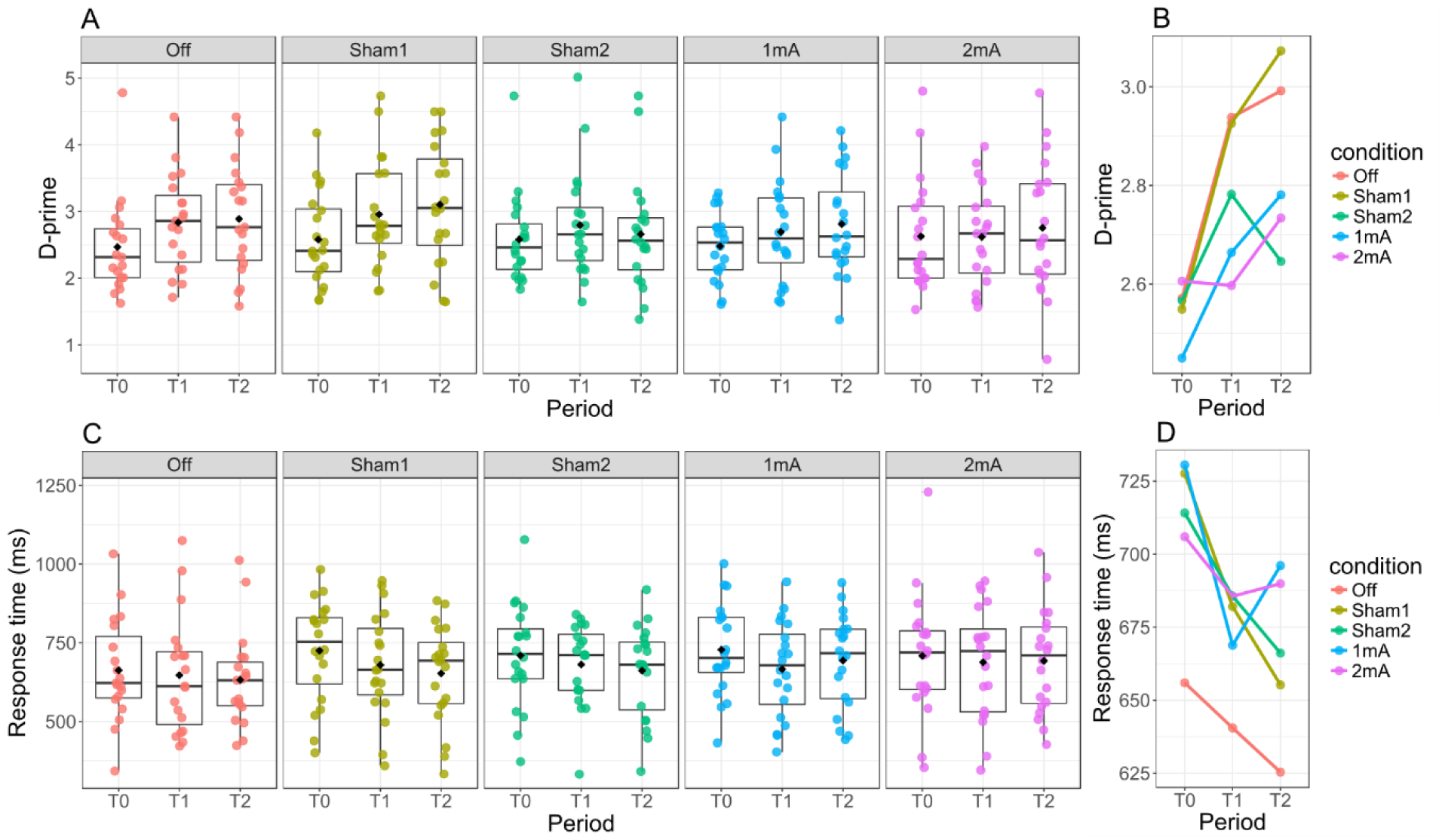
Working memory performance. Individual working memory raw scores are displayed with box plots showing 50% of the data distribution (z-score range: −0.675 to 0.675), centre lines marking median values, and black diamonds representing mean values (panels A and C). Estimated marginal means of working memory scores were obtained from mixed effects repeated measures model analyses (panels B and D). **A)** D-prime scores with individual data. **B)** D-prime scores from estimated marginal means. **C)** Response times for correct responses with individual data. **D)** Response times from estimated marginal means. T_0_: baseline; T_1_: during-tDCS; T_2_: post-tDCS.

There was a significant main effect of Time (*F*_(2,186.9)_ = 6.43, *p* = 0.002) for 3-back response times. Main effects for Baseline Performance (*F*_(2,93.8)_= 2.81, *p* = 0.065) and Condition (*F*_(4,95.3)_ = 0.50, *p* = 0.733) were not significant. The Time × Condition interaction was not significant (*F*_(8,186.9)_ = 0.83, *p* = 0.577); see Figure 3C and 3D for raw scores and plots of estimated marginal means, respectively. Planned simple contrasts for the post-tDCS time point revealed no significant effect of Off compared to Sham1 (*p* = 0.558, Hedges’ *g* = 0.19) or Sham2 (*p* = 0.426, Hedges’ *g* = 0.26).

### Electroencephalography (EEG) event related potentials

Three participants out of the sample of 99 were considered outliers and excluded from all neurophysiological analyses.

Results from the MRMM showed no significant main effect of Time (*F*_(1,54)_ = 0.02, *p* = 0.886) for mean P3 amplitude. Main effect for Condition (*F*_(2,54)_= 3.87, *p* = 0.027) and the Time × Condition interaction (*F*_(2,54)_= 5.42, *p* = 0.007), however, were both significant (see Figure 4). Post-hoc analyses of the Time × Condition interaction effect showed a significant decrease in P3 amplitude from baseline to the post-tDCS time point for Off (*F*_(1,54)_ = 5.04, *p* = 0.029, Hedges’ *g* = −0.27) and a significant increase for 1mA (*F*_(1,54)_ = 5.64, *p* = 0.021, Hedges’ *g* = 0.30). There was no change from baseline to post-tDCS for Sham2 (*F*_(1,54)_ = 0.21, *p* = 0.652, Hedges’ *g* = −0.05).

**Figure 4.**
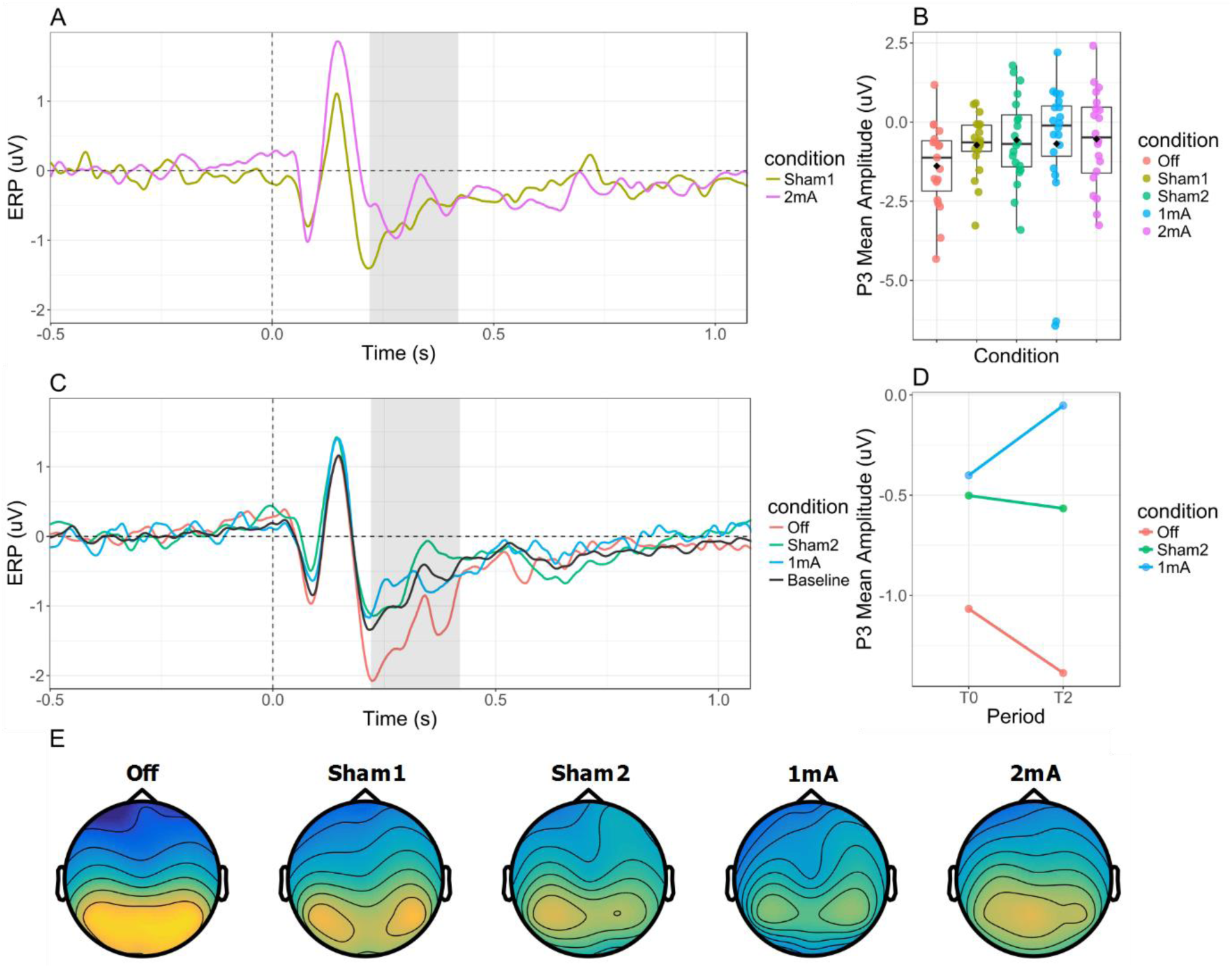
Event-related potentials (ERPs). ERPs were assessed at a midline frontal channel (Fz) following letter presentation (both targets and non-targets) on the 3-back working memory task. **A)** ERPs from Experiment 1 for 2mA and Sham1 tDCS conditions. Shaded regions indicate the time window (220-420ms) used to calculate the mean amplitude of the P3 component. **B)** P3 amplitudes at post-tDCS 3-back task presentation. Individual raw scores are displayed with box plots showing 50% of the data distribution (z-score range: −0.675 to 0.675), centre lines marking median values, and black diamonds representing mean values. **C)** ERPs from Experiment 2 for 1mA, Sham2, and Off conditions. An ERP for the 3-back task at baseline is included using combined data from participants in Off, Sham2 and 1mA conditions. Shaded regions indicate the time window (220-420ms) used to calculate the mean amplitude of the P3 component. **D)** P3 at baseline (T_0_) and post-tDCS (T2) from estimated marginal means. **E)** Spatial topographies of the P3 component for all conditions during the post-tDCS 3-back task.

Analysis of post-tDCS mean P3 amplitudes across the five conditions was significant (F_(4,91)_= 2.77, *p* = 0.032); see Figure 4B for raw scores, including outliers. Post-hoc pairwise comparisons revealed significant differences between Off-2mA (*p* = 0.035, Hedges’ *g* = 0.68), Off-1mA (*p* = 0.002, Hedges’ *g* = 1.07), and Off-Sham2 (*p* = 0.042, Hedges’ *g* = 0.66), but not Off-Sham1 (*p* = 0.052, Hedges’ *g* = 0.64).

### Association between behavioural and electrophysiological measures

Changes in P3 amplitude were correlated with changes in working memory d-prime (r = 0.34, *p* = 0.010) but not response time (r = −0.14, *p* = 0.300) following tDCS compared to baseline levels (Figure 5A-B). Therefore, an increase in P3 mean amplitude is associated with an increase in d-prime.

**Figure 5.**
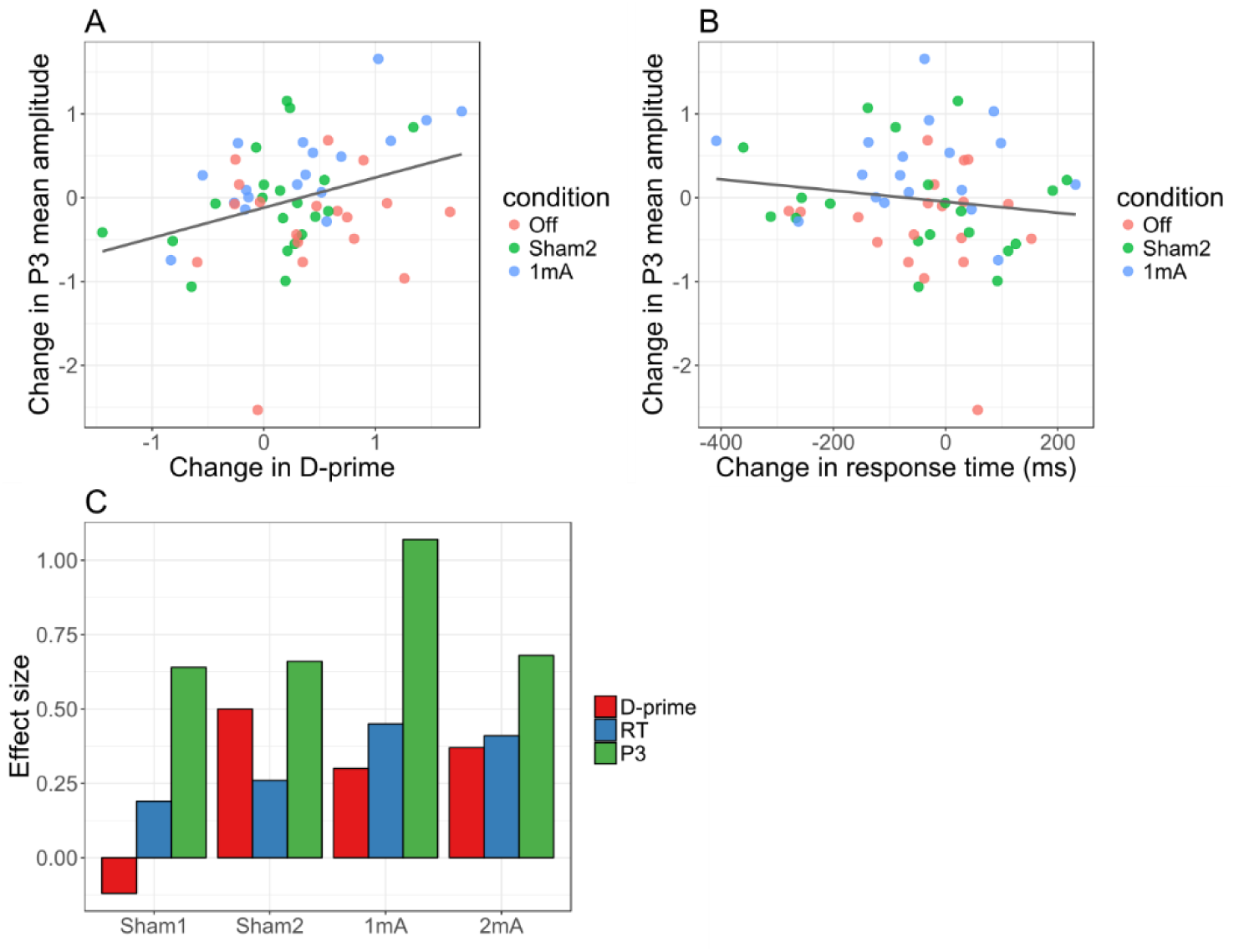
Relationship between working memory performance and event-related potentials. Correlations using change scores, calculated by subtracting baseline from post-tDCS scores, are shown for: **A)** D-prime, a measure of discriminative sensitivity, was significantly correlated with P3 amplitude; **B)** Response time (RT) was not correlated with P3 amplitude. **C)** Effect sizes (Hedges’ *g*) for behavioural and electrophysiological measures comparing conditions against Off at the post-tDCS time point. N.B. The additive inverse of D-prime effect sizes is displayed (i.e. the negative of the value).

Hedges’ *g* statistics, calculated by comparing each condition against the Off condition at the post-tDCS time point, showed that electrophysiological effect sizes were approximately double those for behavioural outcomes (Figure 5C).

### Participant blinding

Pearson chi-square test for participants’ guess of tDCS condition was significant (*p* = 0.043, Cramer’s *V* = 0.32; see Table 1). Post-hoc testing of adjusted standardised residuals showed this significance was due to the Off condition, in which a higher proportion of participants correctly deduced they were receiving sham stimulation. To explore whether differences in participant guesses may explain our findings, we performed additional independent samples t-tests comparing participants who guessed they had received active tDCS (n = 35) against those who guessed they had received sham tDCS (n = 64), regardless of stimulation condition, for each of the behavioural and neurophysiological outcomes at the post-tDCS time point. None of the outcomes assessed were significantly different (working memory response time: *p* = 0.564; working memory D-prime: *p* = 0.397; P3 mean amplitude: *p* = 0.069).

## Discussion

Existing evidence suggests that tDCS can produce cognitive enhancement, however, optimal stimulation parameters, including current intensity, remain unknown. Additionally, it is not yet clear whether commonly used sham tDCS conditions can produce behavioural and neurophysiological effects. In this study, we investigated dosage effects of tDCS using two current intensities and compared two sham tDCS protocols against a sham condition where participants received no stimulation. No significant differences in working memory performance and response time scores were found between stimulation conditions, suggesting that behavioural measures were not affected by tDCS. In contrast, a neurophysiological measure revealed significant differences between receiving no stimulation and 1mA, 2mA, and one of the sham tDCS conditions, indicating that sham stimulation may indeed be biologically active. Effect sizes for these differences were moderate-large across both active and sham tDCS conditions (Hedges’ *g:* 0.64 – 1.07), with the largest effect produced by 1mA stimulation. This suggests a non-linear relationship between tDCS dose and working memory functioning.

Participants showed a general improvement in working memory performance for both response time and d-prime across multiple task presentations, which is attributed to practice effects. However, we were unable to detect any significant performance differences between tDCS conditions. Lack of significance on the 3-back task is in agreement with the findings of Hoy et al. (13), who used current intensities of 1mA and 2mA to the left DLPFC (delivered ‘offline’, prior to task presentation) to investigate the impact of tDCS dosage on working memory performance. The present study tested the same current intensities (1mA and 2mA), but applied stimulation during task presentation (‘online’ stimulation), and used a higher current density, both of which are thought to be beneficial for cognitive enhancement (8, 40-42). We observed a similar lack of behavioural effects, despite modifications to the study protocol (i.e. online stimulation and higher current density) intended to maximize the effects of tDCS on working memory performance.

Meta-analyses of tDCS effects on working memory show significant improvement in response times with stimulation (6-8). Though significant, the benefits of tDCS in contrast to sham stimulation typically are small, with effect sizes of 0.15 of the standardised mean difference (6), and unreliable between participants (43). Our data adds to the previous literature to further suggest that tDCS does not produce substantial improvements in working memory performance in healthy participants (44), regardless of the tDCS dose used. This finding is particularly relevant to the do-it-yourself (DIY) community seeking to improve cognition using tDCS as a form of electro-doping (45, 46).

Neurophysiological responses to tDCS, assessed using the P3 amplitude at channel Fz, differed significantly, with statistical effect sizes (i.e. Hedges’ *g*) approximately double those obtained using working memory measures of performance (ERP effect sizes ranged from 0.64 – 1.07 whereas behavioural measures ranged from −0.12 – 0.50; see Figure 5C). The P3 component is thought to be generated by at least two subcomponents (47). The earlier subcomponent is predominantly produced in frontal structures such as the DLPFC and anterior cingulate cortex and is involved in attention processes (48, 49). The later subcomponent is generated in the parietal cortex and is linked to working memory updating and processing following attentional reallocation of cognitive resources (50). Correlations between working memory performance and the P3 component have previously been observed, including inverse correlations between occipital P3 amplitude and error rates and response times (33), in addition to working memory load (51, 52). Despite lack of significant findings for working memory performance, a positive correlation was observed between neurophysiological outcomes and d-prime. This would suggest that, on average, participants with the largest increase in P3 also showed the greatest improvement in working memory performance.

Interestingly, the P3 amplitude revealed significant differences between the sham stimulation conditions and the Off condition, suggesting that sham tDCS may not be biologically inert. Notably, the same sham tDCS condition (i.e. Sham2) was used in a multi-centre trial of tDCS efficacy in depression, which found greater improvement in mood after weeks of sham stimulation compared to active treatment (10). Together these findings highlight the need to further explore potential effects of sham tDCS protocols as there are several alternative explanations for our findings. Firstly, participants were significantly better at identifying the Off condition as a non-active protocol, indicating compromised blinding. It is therefore possible that the lack of paraesthetic sensations during Off stimulation reduced levels of distraction, allowing participants to engage more with the task. Improved task engagement could have enhanced practice effects, thus limiting the ability to detect significant differences with active tDCS conditions. However, we observed no significant difference in outcomes between participants who guessed they had received sham tDCS and those that guessed they had received active tDCS. Alternatively, there is evidence to suggest that current intensities as low as 0.02mA, applied for several hours, can have beneficial effects on depressive symptomatology (53, 54). Measurement of the output during both sham tDCS conditions used in the present study indicated they delivered background currents of a similar intensity following completion of the ramp down phase (Sham1: 0.016mA; Sham2: 0.034mA). Though these currents are very low, over the duration of the experiment and combined with the ramp up/down phase, they may deliver a sufficient charge to alter neural activity (Sham1: 134.4mC; Sham2: 66.3mC). Currently, some research trials apply low intensity tDCS (<0.5mA) as a sham condition (19, 20), on the basis of computer modelling data which suggests low currents do not alter neural activity (21). Therefore, if the neuromodulatory effects of sham stimulation identified in the present study are confirmed, research within the tDCS field may require new methods of sham blinding to obtain more accurate comparisons against active stimulation. These may include de facto masking, where participants are informed that they will receive active stimulation and that paraesthetic side effects occur only on rare occasions (55, 56), or development of novel sham electrodes that completely shunt current across the skin rather than passing through brain tissue.

Lastly, our results imply that measures of brain activity may be more sensitive markers of neural alterations due to tDCS, as has been noted in past research (32). Tasks examining working memory assess the functioning of several cognitive processes including attention reallocation, encoding, retrieval and updating (57-59). Behavioural outcomes therefore reflect the combined result of these processes. ERP components, however, may act as markers for individual processing stages, such as attention and memory updating in the case of P3 (49, 50). If tDCS selectively modulates one of these stages, then the neurophysiological measures that capture them will be more sensitive to detect the effects of tDCS than aggregate behavioural outcomes. Hence, assessing neurophysiological measures may also help to understand the mechanisms of action of tDCS by identifying the processes that are affected. Future research should incorporate similar measures of neural activity where possible to supplement behavioural outcomes.

## Limitations

The study was conducted in two experiments and thus randomisation was not possible to all five conditions at once, potentially introducing sampling bias to the results. However, the inclusion criteria and method of recruitment were purposefully kept consistent in both experiments, such that participants were similar in demographic and working memory outcomes at baseline. The current study used a parallel group design to prevent practice effects from repeated exposure to the working memory task (which would have required five separate sessions to complete). However, evidence seems to suggest that the inter-individual response to tDCS is highly variable (29), and is best accounted for using a cross-over design (within-subjects), which would have also increased statistical power. Nevertheless, stratification of participants into high, medium, and low performance categories based on baseline d-prime score allowed for better homogenisation between conditions and thus reduced the inter-individual variability introduced by the parallel group study design.

## Conclusion

There was no difference in working memory performance between the five stimulation conditions tested. Therefore, no optimal dosage of current intensity can be recommended within the context of working memory augmentation in healthy participants. Neurophysiological measures showed significant differences between active stimulation conditions and no stimulation, as well as one of the sham conditions and no stimulation. Sham tDCS may therefore be a biologically active intervention, and as such novel forms of participant blinding may be required in tDCS research to allow for comparisons against a true placebo response.

## Acknowledgements

No funding sources were sought to conduct this systematic review. S Nikolin was supported by an Australian Postgraduate Award. Dr Martin was the recipient of a NARSAD Young Investigator Grant. The authors have no conflicts of interest to declare.

## Supplementary Material

**Supplementary Table S1.** Number of participants that experienced an adverse event in each condition. Events are sorted according to overall likelihood of occurrence, with events most likely to occur listed first. Significance of adverse event occurrence was tested using Pearson Chi-Square tests.

| **Adverse event** | **Off**<br>**(n=19)** | **Sham1**<br>**(n=20)** | **Sham2**<br>**(n=20)** | **1mA**<br>**(n=20)** | **2mA**<br>**(n=20)** | **p-value** |
| --- | --- | --- | --- | --- | --- | --- |
| Headache | 4 | 5 | 7 | 8 | 11 | 0.153 |
| Tingling | 2 | 7 | 6 | 6 | 8 | 0.785 |
| Itching | 0 | 3 | 7 | 9 | 7 | 0.814 |
| Burning | 1 | 6 | 4 | 6 | 7 | 0.070 |
| Fatigue | 5 | 6 | 5 | 4 | 4 | 0.184 |
| Blurred Vision | 2 | 3 | 5 | 6 | 2 | 0.667 |
| Skin Redness | 0 | 0 | 0 | 3 | 7 | N/A |
| Pain | 1 | 2 | 1 | 2 | 3 | 0.359 |
| Nausea | 1 | 0 | 1 | 2 | 5 | 0.247 |
| Dizziness | 1 | 1 | 3 | 2 | 2 | 0.273 |
| Pressure | 1 | 1 | 3 | 2 | 0 | 0.133 |

